# Streamlined intravital imaging approach for long-term monitoring of epithelial tissue dynamics on an inverted confocal microscope

**DOI:** 10.1101/2023.05.10.540242

**Authors:** Michael Hamersky, Khushi Tekale, L. Matthew Winfree, Matthew JM Rowan, Lindsey Seldin

## Abstract

Understanding normal and aberrant in vivo cell behaviors is necessary to develop clinical interventions to thwart disease initiation and progression. It is therefore critical to optimize imaging approaches that facilitate the observation of cell dynamics in situ, where tissue structure and composition remain unperturbed. The epidermis is the body’s outermost barrier as well as the source of the most prevalent human cancers, namely cutaneous skin carcinomas. The accessibility of skin tissue presents a unique opportunity to monitor epithelial and dermal cell behaviors in intact animals using noninvasive intravital microscopy. Nevertheless, this sophisticated imaging approach has primarily been achieved using upright multiphoton microscopes, which represents a significant barrier-for-entry for most investigators. In this study, we present a custom-designed 3D-printed microscope stage insert suitable for use with inverted confocal microscopes that streamlines long-term intravital imaging of ear skin in live transgenic mice. We believe this versatile invention, which may be customized to fit the inverted microscope brand and model of choice, as well as adapted to image additional organ systems, will prove invaluable to the greater scientific research community by significantly enhancing the accessibility of intravital microscopy. This technological advancement is critical to bolster our understanding of live cell dynamics in both normal and disease contexts.

**SUMMARY:** A new tool to simplify intravital imaging using inverted confocal microscopy.

## INTRODUCTION

Intravital microscopy is a powerful tool that allows the monitoring of cell behaviors in their unperturbed in vivo environments. This unique method has provided key insights into the inner workings of complex mammalian organ systems, including lung^1^, brain^2^, liver^3^, mammary gland^4^, intestine^5^, and skin^6^. Furthermore, this approach has revealed cell behavioral alterations during tumor development^7^, wound healing^8, 9^, inflammation^10^, and other diverse pathologies in situ. In this study, we focus on enhancing the accessibility of intravital microscopy to image live epithelial and stromal dynamics in intact mouse skin. Understanding cell behaviors in mammalian skin is of high clinical importance due to the remarkable regenerative and tumorigenic capacity of this tissue.

Intravital imaging in mice had been primarily performed using upright multiphoton microscopes due to their ability to provide high resolution imaging at tissue depths > 100 μm^11,12^. Nevertheless, these instruments lack the workhorse versatility and more general accessibility of inverted confocal microscopes, which are more user-friendly, provide the ability to image cultured cells, do not require complete darkness during image acquisition, are generally safer, among other notable advantages. In this study, we present a new tool that significantly enhances intravital imaging accessibility by adapting this approach for inverted confocal microscopes.

Here, we present a 3D-printed custom stage insert design that incorporates several key features to facilitate stable long-term intravital imaging of mouse ear skin on an inverted confocal microscope (Figures 1-5). These specialized features include an offset objective hole that allows the full body of an adult mouse to lay entirely flat during imaging. This minimizes the vibrational interference of mouse body movements on imaging and eliminates the need to administer controlled substances such as ketamine and xylazine to dampen breathing, a practice often coupled with intravital imaging^6^. In addition, corner brackets on the insert correctly position an isoflurane nose cone to align with the face of the mouse, a metal ear clip immobilizes the mouse ear to a custom-built coverslip disk, and an optional detachable closed-loop biofeedback heat plate lies flush within the insert to stabilize mouse body temperature during long imaging sessions. The custom coverslip disk, which provides a flat surface essential for the mouse head and ear to lay flat, was generated in a machine shop by removing the walls of a generic coverslip-containing cell culture dish. The use of a 40x silicone oil immersion lens (1.25 Numerical Aperture (N.A.), 0.3 mm working distance) in conjunction with the coverslip disk and custom stage insert provides high resolution images > 50 μm deep into the ear dermis.

**Figure 1.**
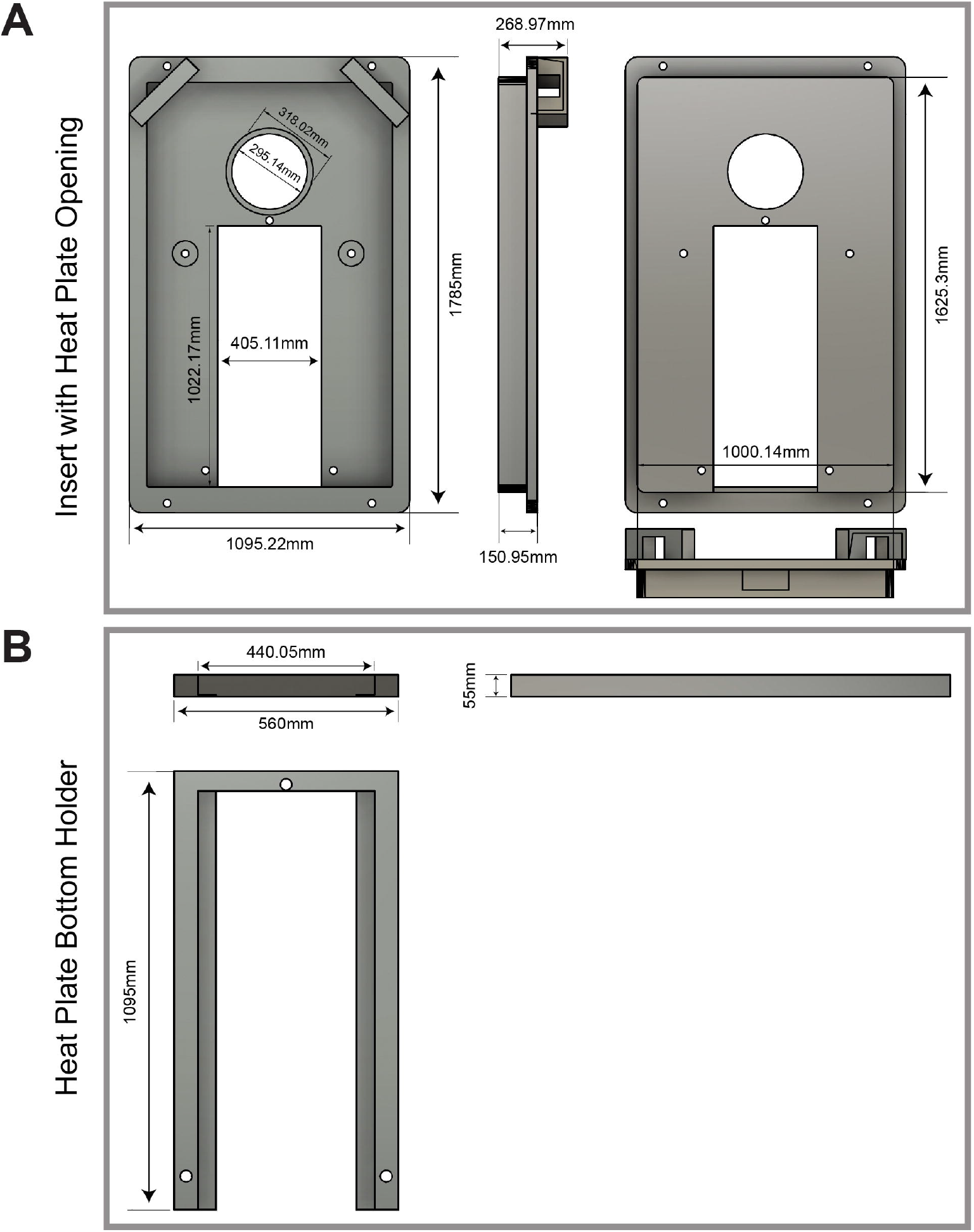
Customized 3D-printed stage insert design. (A-B) Design and dimensions of custom 3D-printed insert with heat plate opening (A), as well as heat plate bottom holder (B), which is printed separately and then screwed into insert.

To test the functionality of this new inverted microscope stage insert, we captured z-stacks spanning all epidermal epithelial layers over a 3-hour time-course in the ear of a live transgenic K14-H2B-mCherry^13^ adult mouse (epithelial nuclei in this mouse line contain a red fluorescent label) (Figure 6A-A’). We also captured z-stacks spanning several fibroblast layers within the skin dermis over a 3-hour time-course in the ear of a live transgenic Pdgfra-rtTA^14^; pTRE-H2B-GFP^**15**^ adult mouse (fibroblast nuclei in this mouse line contain a green fluorescent label following doxycycline induction) (Figure 6B-D’). Our high-resolution data demonstrate consistent stability by lack of drifting in the x, y, and z planes, thus proving the effectiveness of this new intravital imaging tool for use on inverted microscopes. Importantly, the dimensions of this 3D-printed stage insert can be adjusted to fit any inverted microscope, and the positioning of the objective opening can be moved to alternative locations within the insert to better suit imaging a particular tissue and/or animal model of interest. This invention can thus empower individual laboratories, or investigators with core facility confocal access, to adapt this tool for their unique intravital imaging needs, thereby streamlining evaluation of diverse in vivo cell biology.

## PROTOCOL

This research was performed in compliance with Emory University and Atlanta Veterans Affairs Medical Center animal care and use guidelines.

### 1. Installing Live Imaging Insert on Inverted Microscope Stage

1.1 Insert is constructed using .stl files specifying 3D dimensions and design (see Figures 1, 2A), a 3D printer, and polylactic acid (PLA).

**Figure 2.**
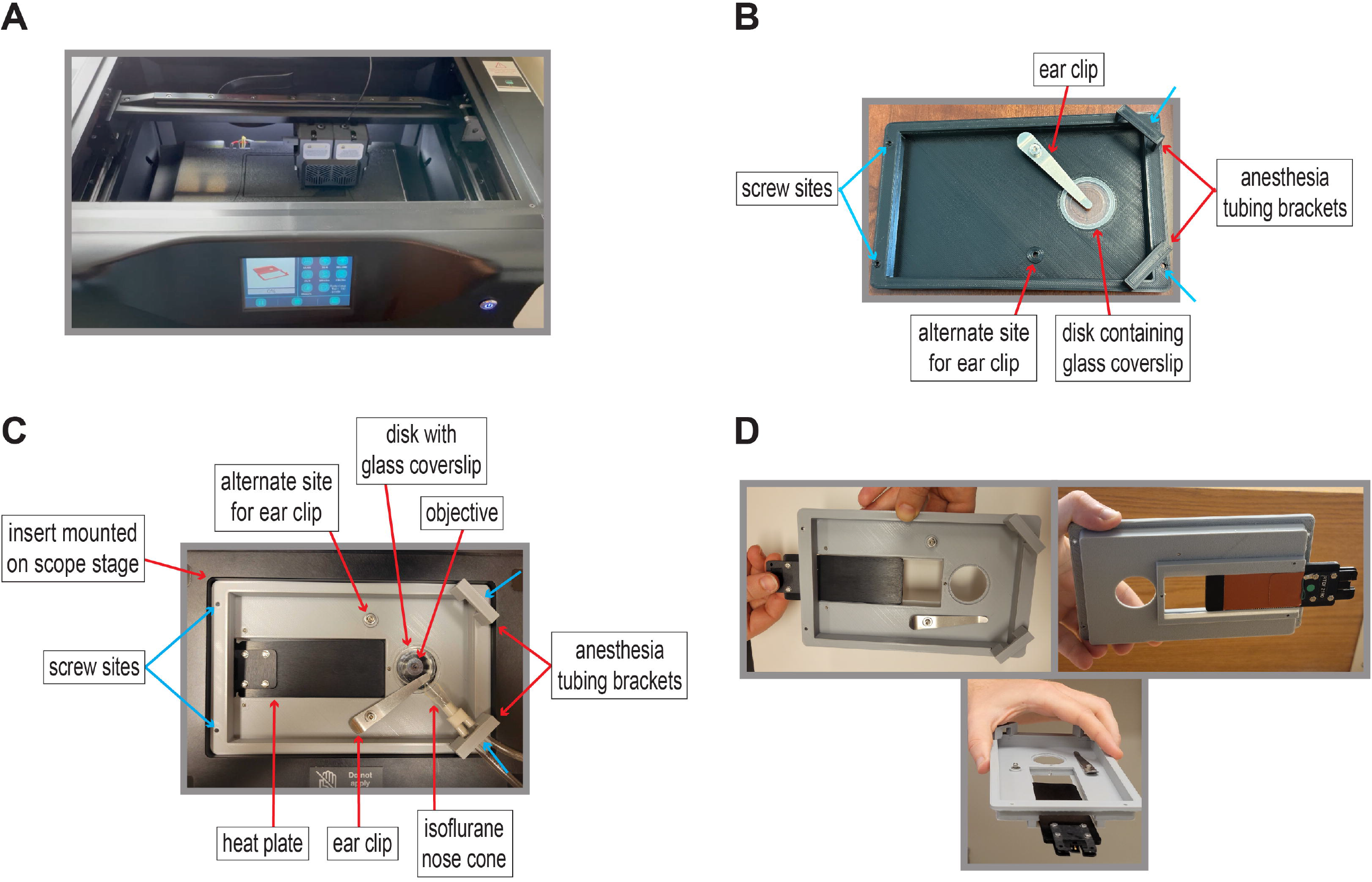
New stage insert streamlines intravital imaging on inverted confocal microscopes. (A) Insert being constructed using a 3D printer. (B) Simple insert model without heating device: Live imaging insert contains four screw sites (blue arrows) for microscope stage attachment. Metal ear clip flattens and immobilizes ear onto 35 mm wide plastic disk containing a 20 mm wide glass coverslip. Insert contains two options for ear clip placement to provide flexibility with mouse orientation. Asymmetric placement of hole allows adult mouse to lay flat. Side brackets align and immobilize isoflurane nose cone to facilitate mouse attachment. Simplified model requires placement of small heating pad (or alternative heat source) under mouse to regulate body temperature. (C) Advanced insert model with built-in heat plate. (D) Heat plate is installed by sliding into grooved opening of insert through a side opening.

1.2 Carefully place insert (Figure 2B-C) into large microscope stage groove (Figures 2C, 3A).

**Figure 3.**
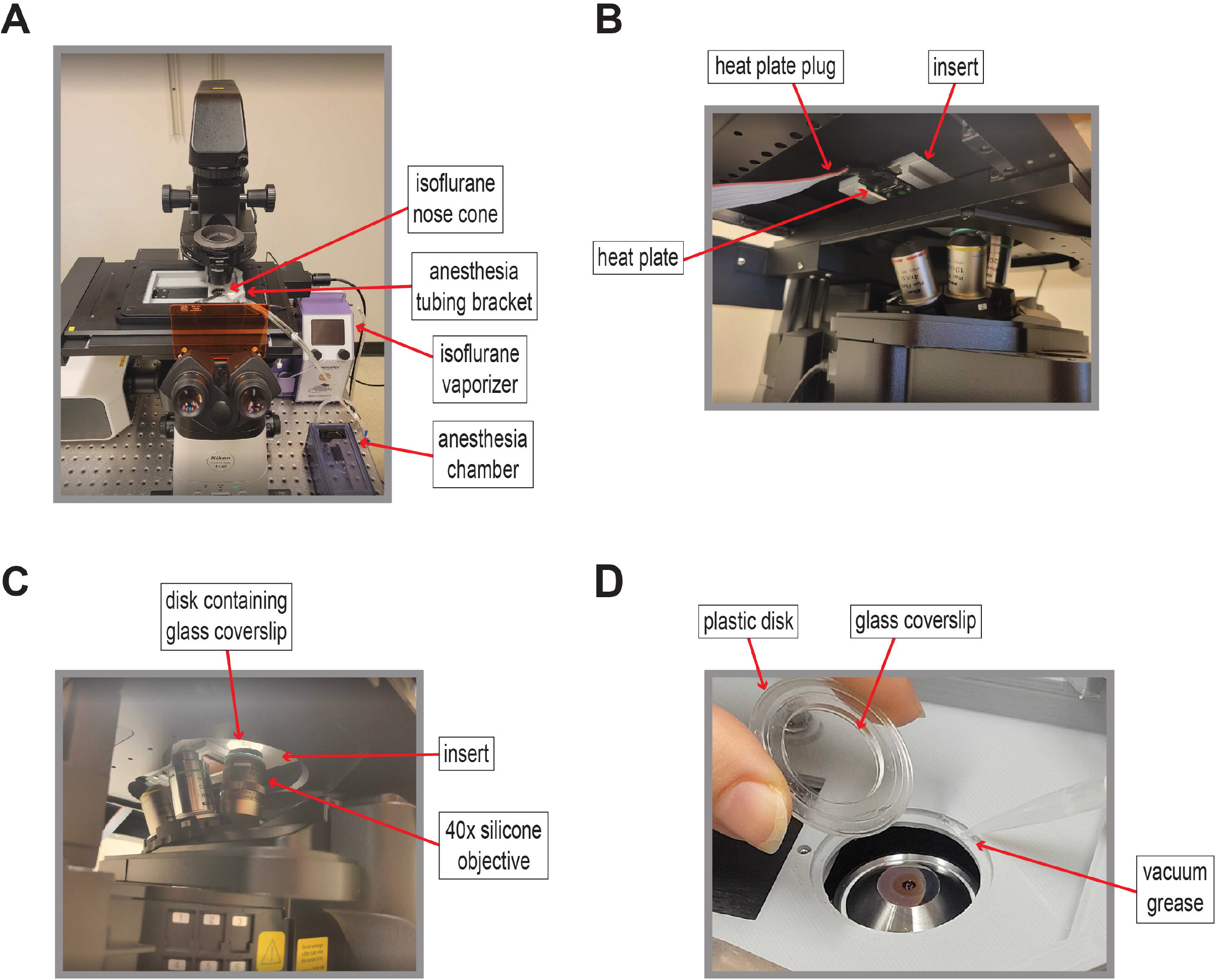
Intravital imaging insert installed on microscope stage. (A) Insert mounted on stage of Nikon AxR laser scanning inverted confocal microscope. Proximity of isoflurane vaporizer and chamber allows threading of nose cone tubing through insert bracket. (B) Heat plate plug extends below microscope stage to connect to controller. (C) Insert hole aligns with 40x silicone objective. (D) Plastic disk (35 mm diameter) containing glass coverslip (20 mm diameter) is laid atop the grooved opening of the stage insert and sealed in place with vacuum grease. Coverslip disk was created by removing walls of 35 x 10 mm glass bottom cell culture dish.

1.3 Use screws to secure insert via four screw holes located in each corner of insert (Figure 2B-C).
  1.3.1 NOTE: Insert is bidirectional and can be rotated 180 degrees depending on orientation of microscope, stage, and anesthesia apparatus.
1.4 Slide heat plate into insert with plug side down (Figure 2D) so the unit lays atop the lower insert grooves with plug port passing underneath the stage (Figure 3B).

1.5 Align grooved circular opening in insert with 40x silicone oil immersion objective (Figure 3C) and apply silicone oil to center of objective.
  1.5.1 NOTE: If imaging over longer time periods (>1 hour), it is important to apply a generous dollop of oil that will persist at the coverslip/objective interface throughout imaging. However, do not allow oil to spill over the edges of the lens.
1.6 Use syringe to apply small quantity of vacuum grease along grooved circular opening and lay coverslip disk atop to seal onto insert (Figure 3D).
  1.6.1 NOTE: Coverslip disk is created by removing walls of 35 x 10 mm glass bottom cell culture dish (performed by machine shop).
1.7 Raise 40x objective until it kisses the bottom of the glass coverslip.

### 2. Isoflurane Configuration and Mouse Prep

2.1 Position low-flow electronic vaporizer components (anesthesia chamber, tubing, vaporizer, charcoal canister) to allow nose cone and attached tubing to reach insert (Figure 3A).
  2.1.1 NOTE: CAUTION! Isoflurane is an inhalation anesthetic and should be handled with care to avoid spills and minimize human exposure.
2.2 Measure weight of isoflurane bottle prior to use and log.
2.3 Attach cap from electronic vaporizer to isoflurane bottle.
2.4 Connect power cord to electronic vaporizer. Close blue nose cone clips and open white induction chamber clips to allow airflow though chamber into charcoal canister.
  2.4.1 NOTE: Remove red ambient air cap from left side of machine to allow air flow.
2.5 Allow the system to warm-up for 5 min at 200 mL/min (Low Flow) and 2 % isoflurane.
  2.5.1 NOTE: Although the nose cone setting is selected, the blue nose cone line should remain closed, and the white induction chamber line should remain open.
2.6 Once system is properly equilibrated, purge lines to remove remaining isoflurane from chamber.
2.7 Place mouse in induction chamber and select “High Flow”.
2.8 When mouse is fully anesthetized, select “High Flow” again to stop isoflurane delivery. Purge chamber before opening and adjust clips (blue open, white closed) to deliver isoflurane through the nose cone. Select “Low Flow” while the nose cone is attached to mouse to continue isoflurane delivery.
  2.8.1 NOTE: Confirm mouse is fully anesthetized using ‘toe pinch reflex’ method. With the leg extended, use fingernail to firmly pinch toe without causing physical damage. If the mouse exhibits a positive reaction to the stimulus (i.e., leg retraction, foot twitch, etc.), continue administering anesthetic within the chamber until no reaction is observed. Once appropriately anesthetized, mouse breathing rates should slow to ∼55-65 breaths per minute^16^.
  2.8.2 NOTE: Mouse prep (i.e., hair removal, topical drug delivery, eye ointment application, etc.) should be completed prior to laying animal’s body atop insert.
2.9 Thread isoflurane tubing with attached nose cone through corner tubing bracket on insert (Figures 3A, 5).

### 3. Mouse Placement on Insert for Intravital Imaging

3.1 Plug heat plate into controller (containing attached anal probe), power on, and allow plate to reach 36 °C (Figure 4A).

**Figure 4.**
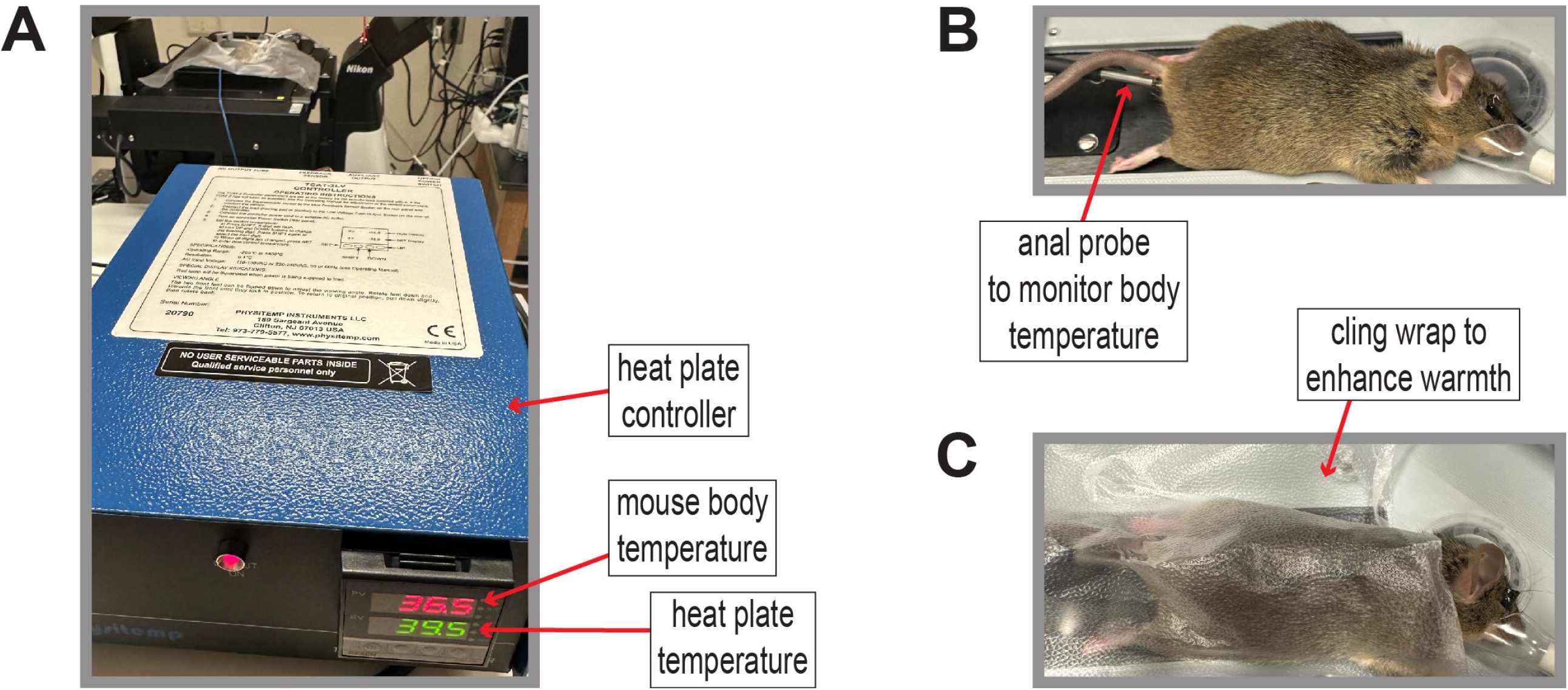
Monitoring animal body temperature using heat plate controller. (A) Heat plate controller, which can be adjusted to stabilize mouse body temperature at the optimal 36 °C throughout intravital imaging session. (B) Anal probe is used to monitor mouse body temperature once mouse is laid atop heat plate. (C) Plastic cling wrap can be used to trap heat to further elevate mouse body temperature.

3.2 Remove anesthetized mouse from induction chamber and lay across heat plate (Figures 4B, 5A).

**Figure 5.**
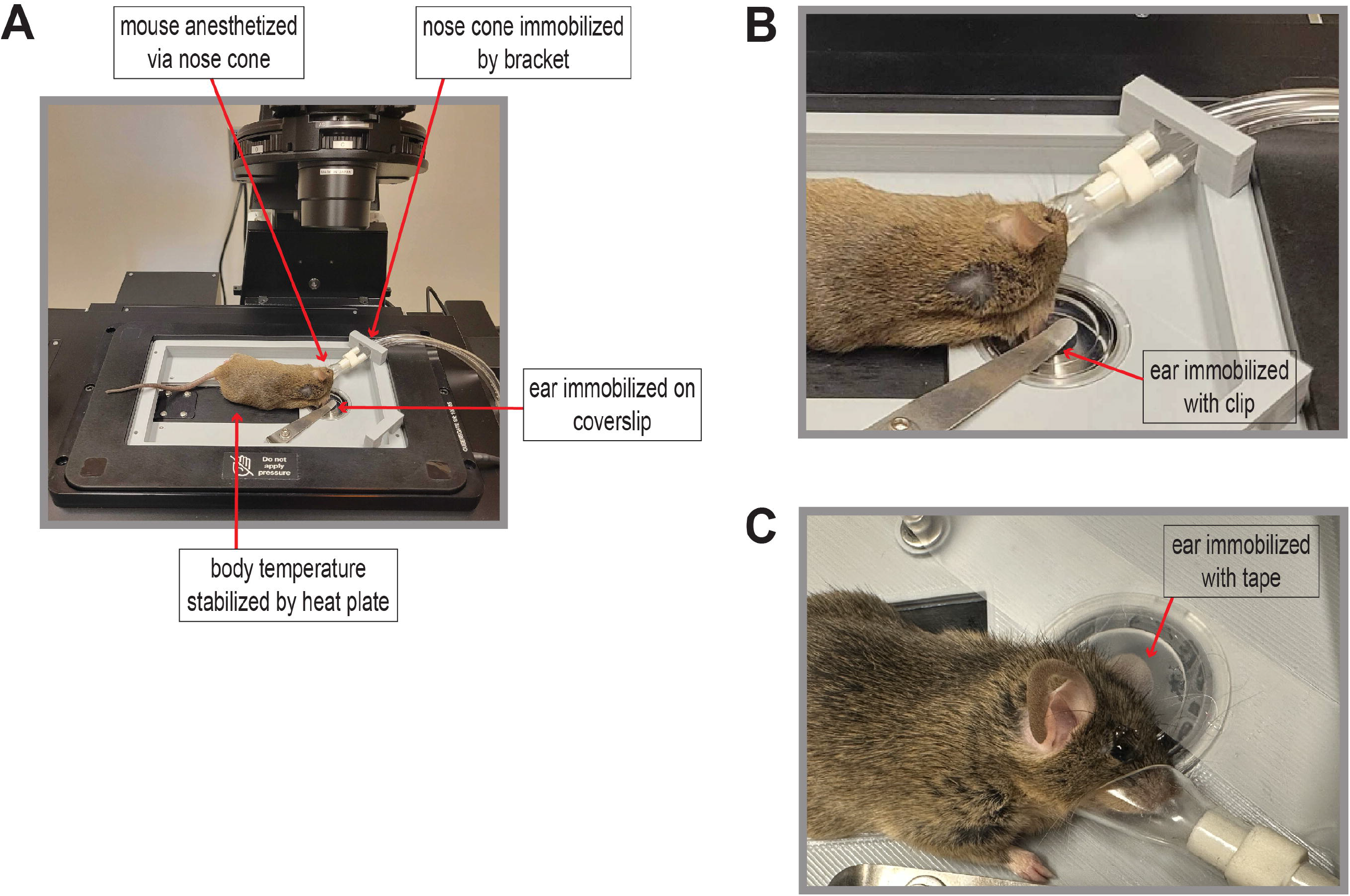
Mouse positioning on intravital imaging insert. (A) Mouse body is spread along top of heat plate with ear centered onto glass coverslip and immobilized with metal ear clip. Bracket positions isoflurane nose cone for mouse attachment. (B) Zoomed-in region of A showing mouse ear immobilization with metal ear clip on glass coverslip and isoflurane nose cone attachment to mouse. (C) Tape can be used as an alternative method of ear immobilization onto glass coverslip.
  3.2.1 NOTE: Transfer from chamber to insert should be rapid to minimize time mouse is without anesthesia.
3.3 Secure nose cone onto mouse (Figures 4B-C, 5).
  3.3.1 NOTE: If necessary, tape can be used to further secure angle and positioning of nose cone.
3.4 Insert anal probe and adjust controller temperature until mouse body temperature is stable at ∼36 °C (Figure 4A-B).
  3.4.1 NOTE: After mouse is properly positioned, plastic cling wrap can be added to trap heat to further elevate body temperature (Figure 4C).
3.5 Position mouse so head aligns with coverslip disk and immobilize ear onto center of glass coverslip using metal ear clip (Figure 5A-B) or tape (Figure 5C).
  3.5.1 NOTE: Pressure of metal ear clip onto ear can be adjusted by loosening the screw that secures the clip to the insert. A metal spring can be added for increased clip tightening flexibility.
  3.5.2 NOTE: To change ear clip location, unscrew the bolt with a 2.5 mm Allen wrench and transfer to secondary site (Figure 2B-C). When reassembling ear clip, clip should be placed against the insert followed by the washer on top. Use bolt to securely fasten clip with slight force to pivot clip.
3.6 Adjust objective z-positioning until cells are within the focal place (Figure 6A, D). Set z-stack and time-lapse parameters according to experimental goals and commence image acquisition.

**Figure 6.**
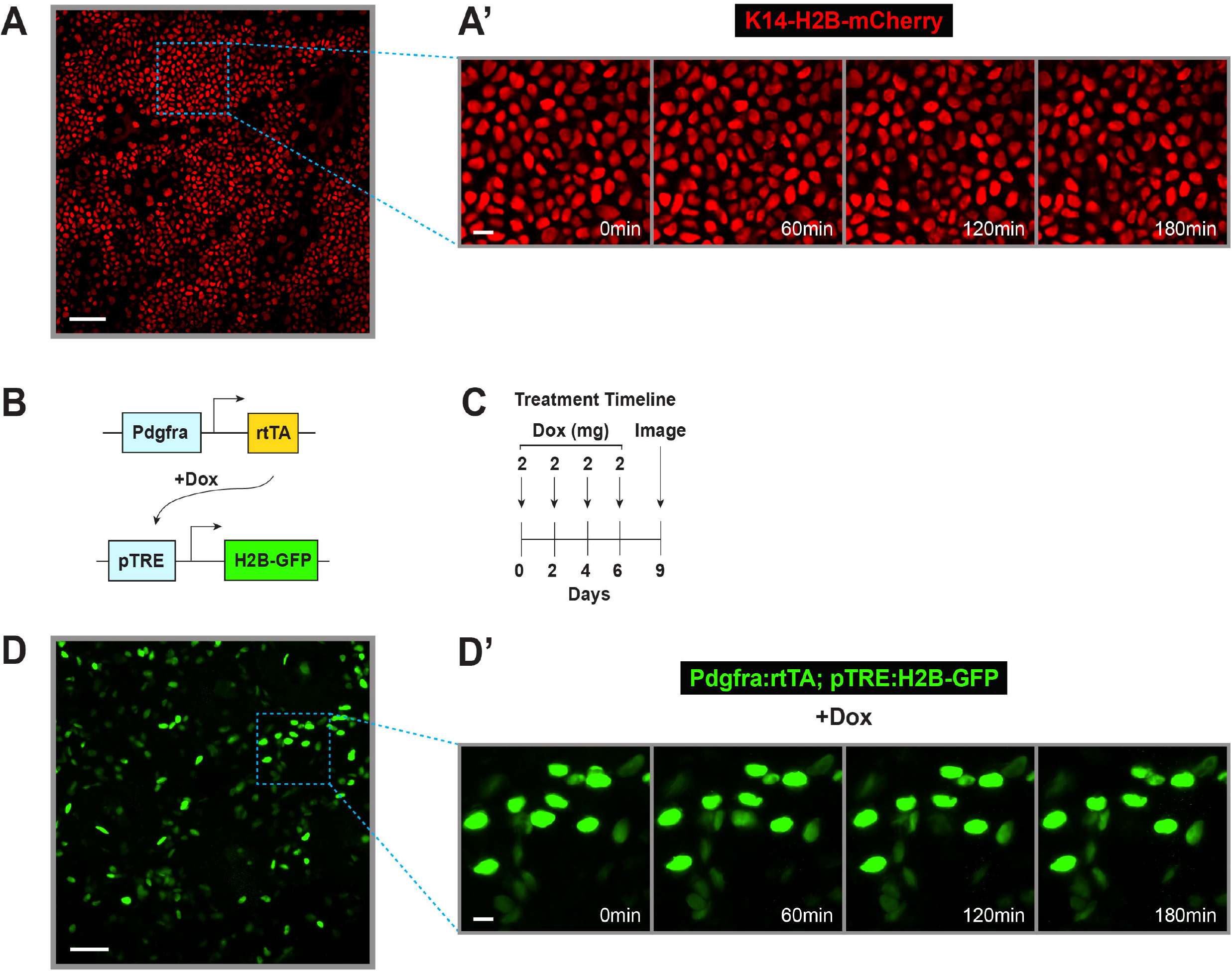
Insert facilitates stable long-term intravital imaging of mouse ear epidermis and fibroblasts. (A) Single z-plane captured from performing intravital imaging on ear epidermal epithelium of 3-month-old adult male K14-H2B-mCherry transgenic mouse. Dotted box indicates zoomed-in region shown in (A’). Scale bar, 50 μm. (A’) Zoomed-in region of (A) showing images every hour over a 3-hour movie. Scale bar, 10 μm. (B) Schematic of doxycycline-inducible transgenic system used to promote in vivo GFP labeling of dermal fibroblast nuclei. (C) Timeline of doxycycline injections. (D) Maximum intensity projection (representing 54 μm total z-depth) of dermal fibroblasts captured by performing intravital imaging on the ear of a dox-injected 8-month-old female Pdgfra:rtTA; pTRE: H2B-GFP transgenic mouse. Dotted box indicates zoomed-in region shown in (D’). Scale bar, 50 μm. (D’) Zoomed-in region of (D) showing images every hour over a 3-hour movie. Scale bar, 10 μm. These time-courses demonstrate stability of long-term intravital imaging using the 3D-printed custom insert.
  3.6.1 NOTE: If correct set-up is achieved, mouse ear can be imaged for at least three consecutive hours (Figure 6B).
  3.6.2 Once z-stack boundaries are set, adjust laser power and gain to ensure no z-plane is oversaturated to minimize photobleaching. Time-lapse parameters are determined by the total thickness of the z-stack, step numbers (Nyquist sampling recommended), and time intervals.
  3.6.3 Imaging parameters (time intervals, total imaging time, etc.) can be determined by multiple factors, including animal viability under anesthesia, laser-induced photobleaching/phototoxicity, and goal of live imaging (i.e., cell division dynamics, cell-cell interactions, etc.).
  3.6.4 Images from four 60-minute time-points displayed in Figure 6A’ were selected from a 3-hour time-lapse movie of mCherry+ epidermal cells in the ear of a 3-month-old adult male K14-H2B-mCherry mouse (∼30 g) captured at 2-minute intervals using a z-step of 0.246 μm to achieve Nyquist sampling across a total z-depth of 24 μm (99 z-stack images acquired per time-point).
  3.6.5 Images from four 60-minute time-points displayed in Figure 6D were selected from a 3-hour time-lapse movie of GFP+ dermal fibroblasts in the ear of an 8-month old adult female Pdgfra:rtTA; pTRE:H2B-GFP mouse (∼30 g) (Figure 6B) captured at 5-minute intervals using a z-step of 2 μm to achieve Nyquist sampling across a total z-depth of 54 μm (28 z-stack images acquired per time-point). This mouse was treated with 2 mg doxycycline every other day for 6 days (4 treatments, 8 mg total) prior to imaging (Figure 6C).

### 4. Termination of Imaging

4.1 Turn on water circulating heat pad and set to “Continuous Cycle” and “35 °C” for at least 30 min prior to termination of image acquisition.
4.2 Upon imaging completion, select “Low Flow” on electronic vaporizer to stop isoflurane delivery and move mouse to heat pad until ambulatory.
4.3 Once awake, move mouse back to transfer container while continuing to maintain on heat pad until fully ambulatory.
  4.3.1 NOTE: Mouse should be consistently monitored while on heat pad until responsive.
4.4 On electronic vaporizer, touch “Select Menu → Anesthetic Control → Empty”. Remove isoflurane bottle from system and replace manufacturer’s cap onto bottle.
4.5 Measure weight of isoflurane bottle and charcoal canister and enter weights into log.
  4.5.1 NOTE: Once charcoal canister weighs 50 g, dispose canister, and replace with new unit.
4.6 Return isoflurane to lockbox.
4.7 Remove coverslip disk and wipe clean with lens paper and lens solution. Store properly to avoid scratches for reuse.
4.8 Remove anesthetic tubing from insert bracket and unscrew insert from microscope stage.

## REPRESENTATIVE RESULTS

Proper assembly of the live imaging insert on an inverted confocal microscope and appropriate orientation of a transgenic mouse atop the insert will be validated by acquiring z-stacks of fluorescently-labeled live ear tissue over a time-course of one or more hours with minimal evidence of drift in the x, y, and z axes. Images should be captured at consistent intervals (interval time will depend on the biological question, strength of fluorescent signal, etc.) so that cell dynamics and image drift can be tracked over time. Throughout the time-course, monitoring individual z plans to ensure they remain in focus will reveal whether animal movement is interfering with imaging stability. An example of a single z-plan remaining in focus over an extended time-course using the live imaging insert is depicted in Figure 6.

## DISCUSSION

In this study, we present a new tool that facilitates stable, long-term intravital imaging of intact mouse skin epithelia on inverted confocal microscopes. This invention is made of polylactic acid (PLA), which is the most common and inexpensive 3D printable material; all in-house 3D printing costs for this insert amount to < $5. The two separate insert pieces (Figure 1) can be easily assembled using Thorlabs set screws (see Materials File). Notably, the 3D printer files can also be used to order this insert by commercial means. An additional option is to use a computer numerical control (CNC) machine to generate the insert out of anodized aluminum, however this is significantly more costly.

To ensure reliable and efficient imaging over a duration of time using this new tool, it is critical to properly size the live imaging insert according to the user’s microscope stage. The insert design files can be adapted to reflect the appropriate insert dimensions that are compatible with the microscope of choice prior to 3D printing. Accurate insert dimensions with an immobilized coverslip disk will minimize positional (x/y) and focal (z) drift throughout each imaging session. It is also important to ensure that the insert depth is sufficient to pass the heat plate plug port underneath the stage.

Prior to imaging, it is essential to confirm that the mouse is fully anesthetized, its body temperature remains stable ∼36 °C using the anal probe, and the ear is firmly immobilized to avoid movement due to breathing. When using the metal ear clip to secure the ear to the coverslip, prevent cutting off normal blood circulation by ensuring that the ear clamp is not screwed down too tight. It is also crucial to monitor and replenish eye ointment on the mouse when necessary to maintain ocular moisture during long imaging sessions.

While this unique insert design provides a new approach for intravital imaging, it has a few notable limitations. Due to the insert’s low positioning in the microscope stage, stage movement in x and y should be very limited after the objective is positioned and the coverslip disk is sealed into place. This limited movement is critical to avoid damage to the objective. Furthermore, it should be noted that the glass coverslip disk is not currently commercially available. To recreate the coverslip disk used in this study, collaboration with a machine shop may be required.

While we demonstrate that this confocal-based method can achieve stable long-term imaging deep into intact tissue, it should be noted that multiphoton microscopes cause less photodamage and penetrate to greater depths compared with confocal instruments^17^. Therefore, the utility of this tool will rely on the strength of the fluorescent signal in the transgenic animal model of choice as well as the tissue depth required to achieve experimental goals. We therefore recommend using the lowest laser power possible to minimize photobleaching while achieving the desired resolution. It is also important that the objective lens used with this insert has a high N.A. for optimal resolution as well as a long working distance so it can reach the coverslip disk. It should be noted that this approach is not meant to replace the well-validated upright multiphoton-based intravital imaging method as described in Pineda *et al*., 2015^6^. Instead, this new tool is intended to provide an effective alternative to labs that do not have access to multiphoton equipment, prefer a less cumbersome system, and/or own an inverted microscope.

Our new tool is remarkably customizable to individualized experimental requirements and preferences. This insert can be used with or without a heat plate, as some alternative options include the placement of a thin heating pad underneath the mouse, wrapping an objective lens warmer around its torso, and/or trapping heat by covering the mouse with plastic cling wrap (Figure 4C). Furthermore, tape may be used in conjunction with or in place of the ear clip to provide optimal tissue immobilization (Figure 5C). The orientation of the stage insert is reversible, wherein the user can decide whether it’s optimal to orient the objective hole on the right or left side. Furthermore, the insert has built-in alternative placement options for the ear clip and isoflurane tubing to provide maximal flexibility based on desired mouse orientation (imaging the right versus left ear), location of isoflurane set-up, and specific microscope configurations (Figure 2C). The insert is easily installable and removable, with a simplified design intended to be highly user-friendly so that intravital imaging can be made approachable to even novice microscopists. Additionally, the asymmetrically-localized objective hole provides the capacity to image diverse animal models as well as organs of varying sizes.

We intend for this invention to enhance the application of intravital microscopy within individual laboratories as well as microcopy cores that contain inverted confocal instruments. This highly accessible and customizable tool will provide investigators the freedom to visualize live cell dynamics across diverse organ systems to reveal significant cell biological insights.

## ACKNOWLEDGMENTS

We thank Valentina Greco for the K14-mCherry-H2B mice. We are grateful to the Emory University Physics Department Machine Shop for generating the glass coverslip disks. This work was funded by Career Development Award #IK2 BX005370 from the US Department of Veterans Affairs BLRD Service to LS, NIH Awards RF1-AG079269 and R56-AG072473 to MJMR, and I3 Emory SOM/GT Computational and Data Analysis Award to MJMR.

